# Can a Tissue-derived Progression Signature Accurately Predict Colorectal Cancer Stage Transitions in Blood?

**DOI:** 10.64898/2026.06.23.734006

**Authors:** Pritom Sarkar, Prapthi Sarkar

**Affiliations:** Department of Computer Science and Engineering, Stamford University Bangladesh, Dhaka 1217, Bangladesh; Department of Biotechnology and Genetic Engineering, Noakhali Science and Technology University (NSTU), Noakhali 3814, Bangladesh

## Abstract

Colorectal cancer (CRC) is challenging to track because its molecular changes are very complex as the disease progresses, creating significant challenges for robust biomarker discovery. In this study, we developed a machine learning framework by integrating monotonic progression and the StepMiner approach. We conducted external validation to identify reproducible, consistent transcriptomic biomarkers associated with CRC progression. Gene expression datasets were analyzed across four disease states from publicly available GEO: normal colon, adenoma, primary colorectal cancer, and metastasis. First, we identified genes with monotonic expression, then used the StepMiner approach to identify genes that act as ’switches’ between stages. A balanced 74-gene signature was used for machine-learning classification with a Random Forest. External validation showed strong performance in tissue-based datasets. However, tissue-derived signatures and plasma and blood-based datasets showed poor performance, highlighting biological differences between transcriptomic profiles. Cross-filtering between tissue-derived genes and blood expression datasets was performed, which resulted in the selection of 62 blood-compatible gene signatures. Leakage-free retraining on GSE164191 achieved a mean AUC of 0.868 with balanced precision. Functional enrichment analysis showed that these genes are highly active in cancer growth. Specifically, genes CBX3, S100A11, PDK4, NCOR1, and SOX4 demonstrated stable and reliable performance across the validation fold.

Overall, our study presents a progression-aware transcriptomic framework for CRC biomarker discovery and demonstrates the importance of external validation. Additionally, we evaluate whether tissue-derived signatures can predict blood profiles. This proposed approach may help the future development of tissue-based diagnostics and minimally liquid-biopsy strategies for CRC.

To ensure reproducibility, our proposed workflow was automated as a Nextflow pipeline. The tissue-derived model was deployed as an application utilizing Angular, ASP.NET Core, and Plumber (R).

## 1 Introduction

Colorectal cancer (CRC) is one of the most common cancers. It is ranked as the top cause of cancer deaths worldwide [1,2]. While CRC is most common in developed nations, cases are rising rapidly in developing nations due to diet and lifestyle changes. This is creating a huge gap as many countries cannot keep up with the rising number of patients affected by the disease. By 2030, the global burden of colorectal cancer is expected to jump by 60%, and can cause over 2.2 million new cases and 1.1 million deaths worldwide[3,4].

Scientists can now use AI and machine learning to find new biomarkers much faster by using online free databases like Gene Expression Omnibus (GEO); also, high-throughput transcriptomic technologies are publicly available (microarrays, RNA-seq, and other next-generation sequencing platforms)[5]. Multiple studies now integrate GEO and TCGA datasets with machine learning to identify diagnostic gene signatures. Random Forest, SVM, and ANN algorithms are commonly used for analysis and frequently achieve high classification performance (AUC >0.90–0.95) for distinguishing tumor from normal tissue[6–8]. In addition to tissue, researchers are also finding blood and plasma-based transcriptomic biomarkers. They are also developing diagnostic panels that can analyze blood-based molecules, such as mRNA, microRNA, and other circulating transcripts, to detect early colorectal cancer[9,10].

Despite these advances, many limitations persist in existing studies of colorectal cancer (CRC) biomarkers. Many published models rely only on single-cohort datasets, frequently lack independent validation, or utilize high-level feature selection strategies that risk overfitting or information leakage[11,12]. However, external validation of machine learning models and multi-omics-based predictors often reduces performance in independent cohorts, and that reflects issues of cohort heterogeneity, overfitting, and differences in sample types[13,14]. It is hard to get the same results in different studies because everyone uses a different data processing pipeline for handling and analyzing tissue and liquid biopsy samples to collect, test, and look at the data[15,16]. Very few researchers have also tried to use tissue-derived signatures in blood or plasma samples. Translating a progression signature from tissue to blood is still rare.

To address these limitations, we developed a machine learning model inspired by the StepMiner approach to classify colorectal cancer using independent GEO datasets[17]. Initially, monotonic transition-associated genes were identified from the GSE41258 dataset, and subsequently, the exact location of the transition was determined using the StepMiner approach. Several approaches, such as WGCNA and LIMMA, have been widely used to identify hub genes and stage-specific signatures in colorectal cancer[18]. But these methods mainly rely on pairwise comparisons and do not explicitly model the ordered, stepwise progression of tumor advancement. With this in mind, we designed our model.

A balanced feature selection strategy was applied by selecting top-ranked genes from multiple biological transition groups to minimize bias toward highly dominant expression patterns. Next, we trained a Random Forest model and tested it externally validated across independent tissue, plasma, and blood (GSE110223, GSE14333, GSE164191) GEO datasets. We filtered the tissue-derived gene signature in the blood dataset to identify reliable biomarkers that can be used to detect colorectal cancer in blood.

Furthermore, we used GO [19] and KEGG [20] analyses to explore biological rules. Overall, we ensure our method works accurately for different types of patient samples.

Finally, we address the common challenges of bioinformatics reproducibility and user adoption. First, our entire study design is automated via a scalable Nextflow pipeline. Second, we developed a production-ready, full-stack application using Angular, ASP.NET Core, and Plumber.

## 2 Methods

### 2.1 Data Acquisition and Preprocessing

#### Discovery Dataset

We obtained gene expression data by downloading from the GEO database under accession number GSE41258 [21] for different stages of disease progression[22,23]. This included 390 samples. The GEO series matrix file was processed with R using GEOquery. Samples were classified into four distinct stages, and metadata was extracted from the characteristics_ch1 and title fields.

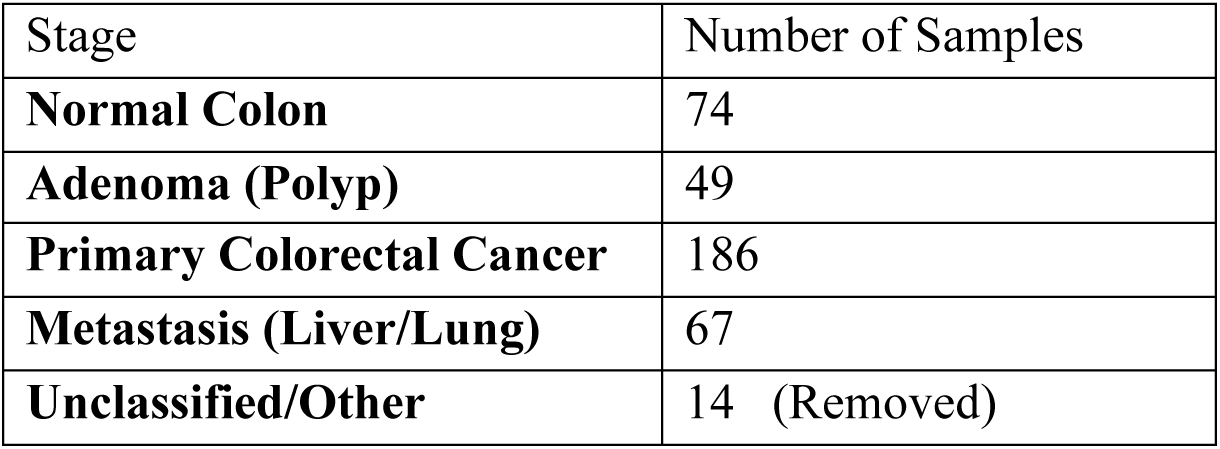

### Data Preprocessing

We processed our discovery dataset (GSE41258) using R pipelines. External validation was performed primarily in Python, except for one dataset, which was validated in R. Gene symbols were converted to hgu133plus2.dbAnnotation, an R package. For genes with multiple probes, we retained the highest absolute logFC value. Probes without a valid symbol were not taken. To contain Ensembl gene identifiers, Ensembl-to-gene symbol mapping was performed before preprocessing. Also, to verify consistency during Python retraining, probe mapping was performed via NCBI annotation guidelines.

### 2.2 Feature Engineering

#### Differential Expression Analysis With LIMMA

LIMMA performed the differential analysis. Four Samples were categorized into a group representing the colorectal cancer progression continuum: Normal Colon, Adenoma, Primary Colorectal Cancer, and Metastasis. We considered the low expression value: any gene entry with an average expression below 5 was excluded. Also, log2 transformation is applied. A design matrix was constructed using the disease stage for experimental purposes, and linear models were fitted for each probe using the limma framework. Pairwise differential expression analyses were performed between consecutive disease stages.

- Normal Colon vs Adenoma
- Adenoma vs Primary Colorectal Cancer
- Primary Colorectal Cancer vs Metastasis

Probes satisfying the criteria of adjusted P-value < 0.05 and absolute log2 fold change (|logFC|) 1 were considered significantly differentially expressed and retained.

#### Progression Gene Identification

To identify monotonic gene expression and to drive the transition between colorectal cancer stages, we also did a progression analysis[21]. The whole algorithm works as follows:

1. To find out the stage (Normal → Adenoma → Cancer → Metastasis), we loaded the DEG results we have. Samples were ordered according to the biological progression sequence.
2. For each candidate gene, we calculated the mean expression per stage and checked if gene expression is monotonic or not. We kept only monotonic genes.
3. For each possible step position i, we calculated the mean of the first segment (position 1 to i) and the second segment (position i+1 to n). For each gene, all possible split positions were evaluated across the ordered samples. At each division position, the mean expression of both groups was calculated, and the sum of squared errors (SSE) was computed. The best split was the one that minimized SSE, indicating the prominent expression transition.
4. To quantify step significance, we calculated F-statistics by following the rule implemented in the original StepMiner algorithm:

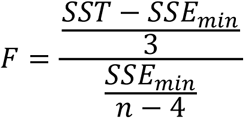

SST indicates the total sum of squares, SSE_min is the lowest error from the best split, and n is the total number of samples. We retained the progression-associated gene where the **F-statistic > 10.**

To find out the biological meaning of those shifts, we matched each gene’s best split point to a specific boundary between cancer stages. Samples were arranged according to progression: Normal Colon → Adenoma → Primary Cancer → Metastasis. Each stage boundary was set at its very last sample. Based on where its split point fell, each gene was assigned to one of three transition groups: Normal-to-Adenoma, Adenoma-to-Cancer, or Cancer-to-Metastasis. This showed us exactly which stage triggered each gene’s major shift in activity and revealed the main genetic changes that occur as each gene progresses.

### 2.3 Stage Classification Model

#### Gene Selection

The top 25 genes were selected for each progression stage (Normal→Adenoma, Adenoma→CRC, CRC→Metastasis) to ensure an unbiased representation of each stage. Eliminates duplicates and left us with 74 unique genes (ranked by F-statistic).

#### Model Training

We implemented a Random Forest Classifier model with 500 trees [25]. To evaluate performance, the dataset was split into 70% for training and 30% for testing. After initial training and validation in R, the model was replicated and retrained in Python for extensive external validation against many independent GEOs. In Python, Random Forest classifiers were implemented using the scikit-learn library and the randomForest package in R[25,26].

#### External Validation of Stage Classifier

To evaluate generalizability and efficiency, we trained 74 genes in GSE41258 (Colon Tissue) [21] and tested on GSE164191 (Colon Blood)[27], GSE110223 (Colon Tissue) [28], GSE142987 (Liver Plasma) [29], and GSE14333 (Colon Tissue - Cancer Only)[30]. Using the R-based pipeline, the model was trained on the GSE41258 [21] dataset and tested using the GSE14333 [30]. Using a Python-based pipeline, the model was retrained on the primary dataset and externally validated on colon blood, colon tissue, and liver plasma.

#### Cross Filtering

Tissue-derived genes were filtered against a blood dataset (GSE164191) [27], and a 62-gene blood-compatible panel was found. We used five-fold cross-validation [31] with a leakage-free feature selection using the Random Forest model [25,32]. Here, we used Python to perform cross-filtering.

### 2.4 Study Workflow

We executed the experiments according to the workflow illustrated in Scheme 1.

**Scheme 1:**
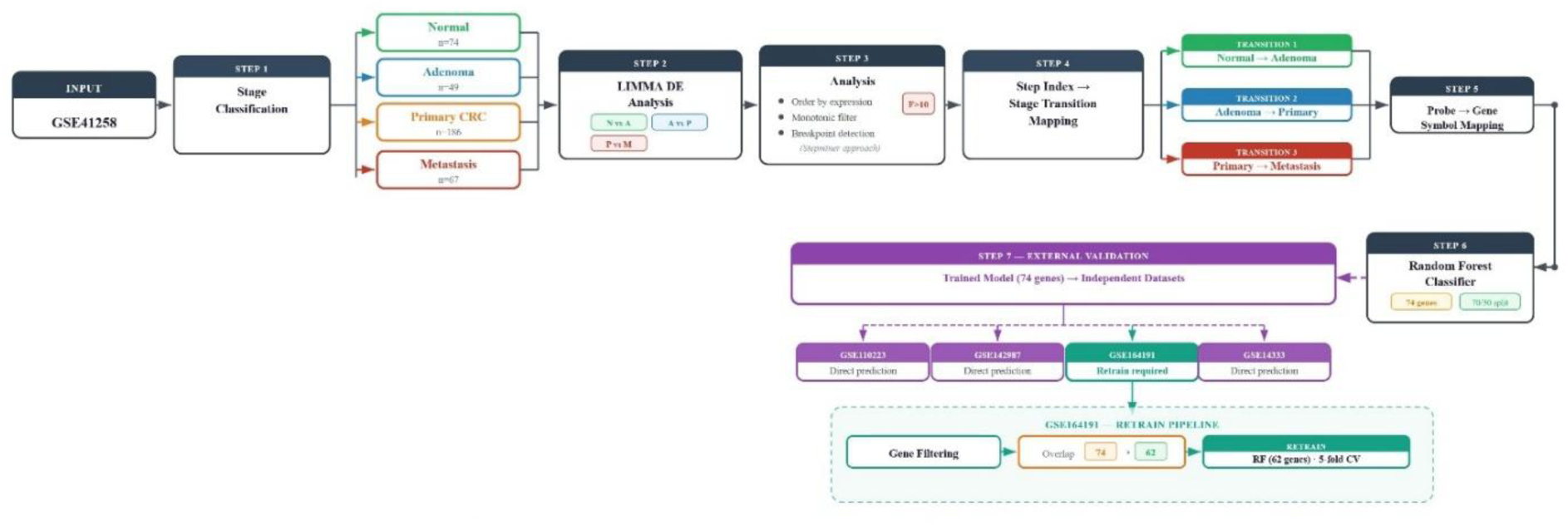
Proposed study design and computational pipeline.

### 2.5 Software Architecture and Workflow

To ensure reproducibility, we automated and containerized all steps into a portable Nextflow pipeline that executes the entire codebase and regenerates all study findings and figures. The finalized tissue model was exported as an .rds file. An application we have built where an R Plumber API takes this saved model and sends its predictions to an ASP.NET Core backend, which displays the results on an Angular frontend.

## 3 Result

### 3.1 Cohort Stratification and Stage-wise Classification

A total of 390 samples from the GSE41258 cohort were classified into biologically relevant colorectal disease stages based on metadata annotations. GEO (GSE41258) included 74 normal colon samples, 186 colorectal cancer samples, 49 adenoma/polyps, and 67 metastatic lesions. Fourteen samples remained unclassified due to ambiguous metadata, and those were excluded. As shown in Figure 1, we have more samples from primary colorectal cancer, which helps us reliably detect gene expression changes across disease stages. While the other samples provide the full disease scenario.

**Figure 1:**
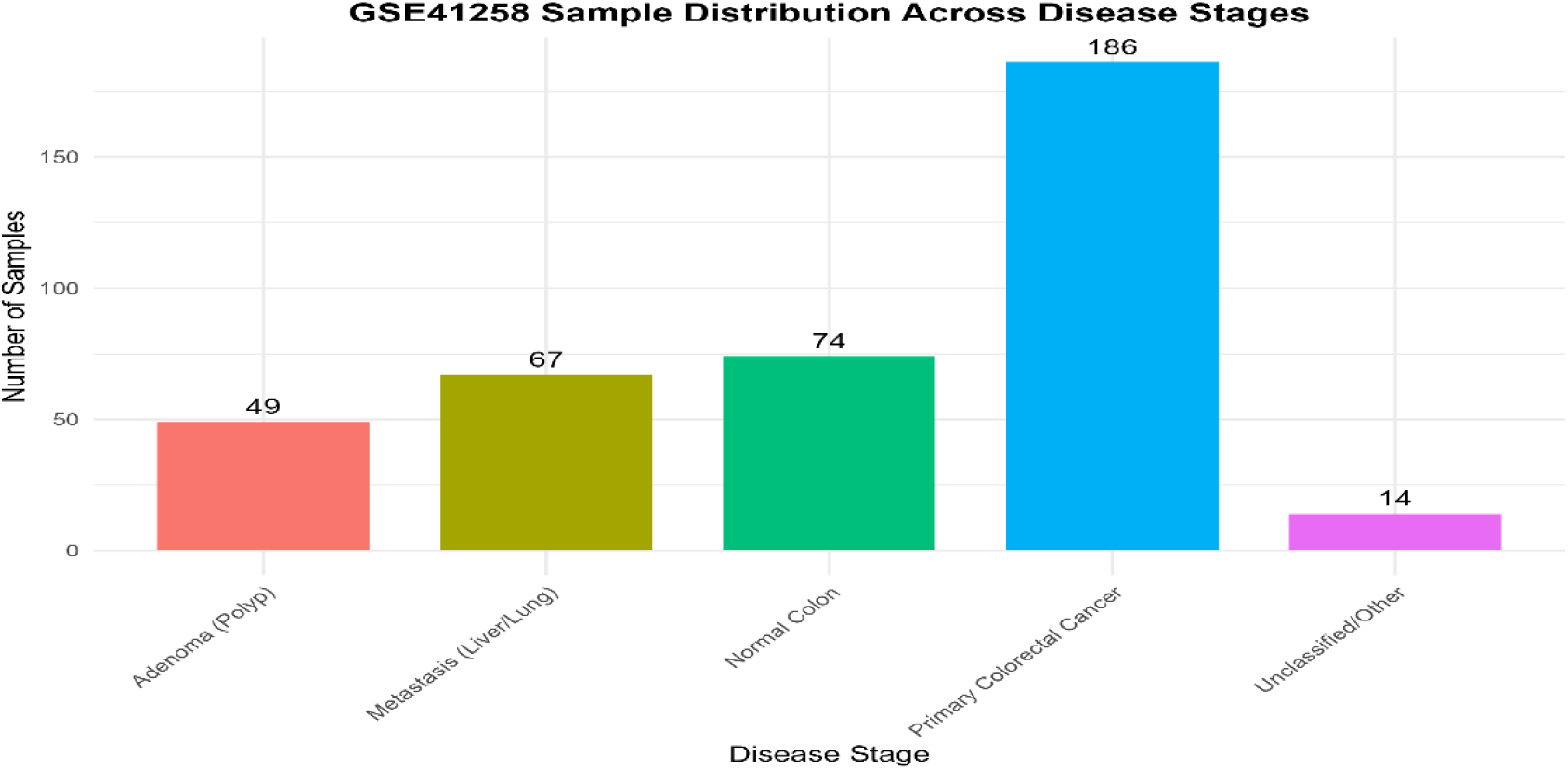
Stage-wise distribution of GSE41258 samples.

### 3.2 Differential expression across disease stages

We used LIMMA to perform pairwise comparisons using this: (log₂FC| > 1, adj.P < 0.05). As Table 1 shows, we found normal-to-adenoma transition identified 1,051 DEGs, the adenoma-to-carcinoma transition identified 377 DEGs, and the cancer-to-metastasis transition identified 136

**Table 1:**
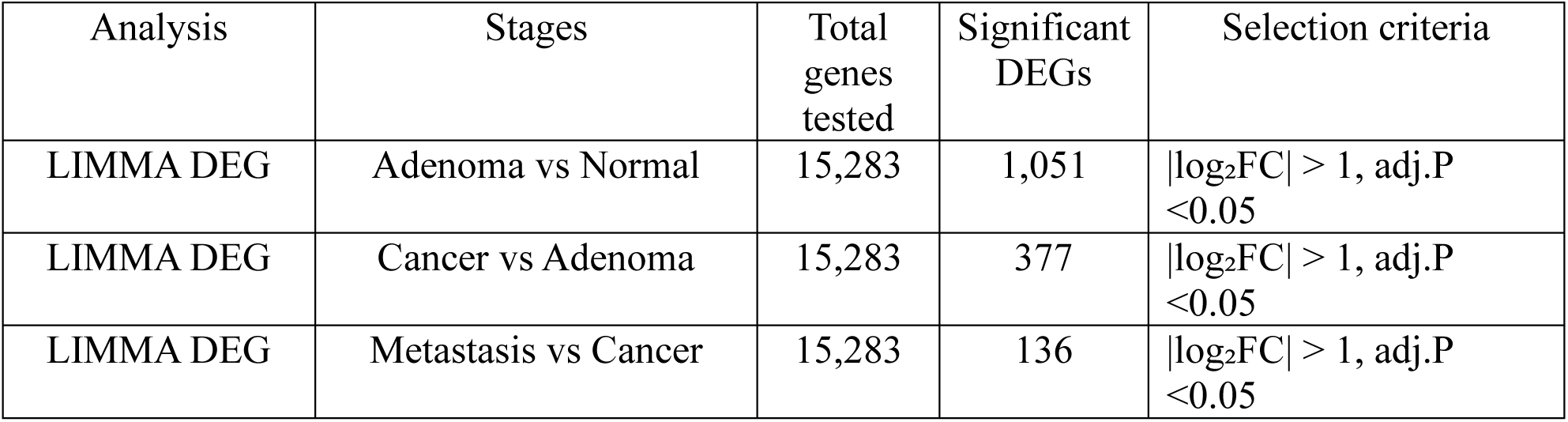
Summary of differentially expressed and progression-associated genes.

DEGs out of a total of 15,283 genes. Figure 2 shows plots of gene expression changes across different stages of cancer (Adenoma vs Normal, Cancer vs Adenoma, and Metastasis vs Cancer). Red dots indicate upregulated genes and blue dots indicate downregulated genes. Genes were taken using the above formula.

**Figure 2:**
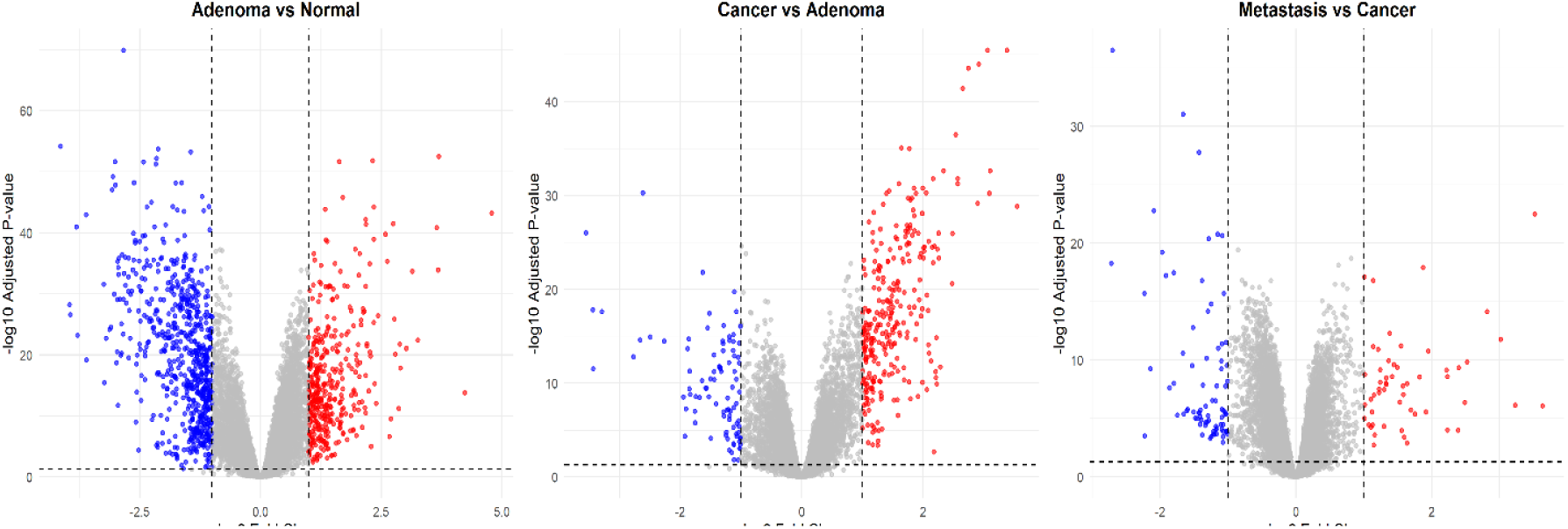
Panel A: Adenoma vs Normal | Panel B: Cancer vs Adenoma | Panel C: Metastasis vs Cancer.

### 3.3 Stepwise Transcriptional Changes Across Stages

To find out progressive gene expression changes across the Normal Colon → Adenoma → Primary Cancer → Metastasis, we performed an analysis inspired by the StepMiner approach on monotonic genes (genes showing consistently increasing or decreasing expression across stages). Among 2,721 monotonic genes, 2,123 genes were found significant stepwise pattern.

F-statistics > 10 were considered significant Genes (Table 2). To biologically transcriptional shifts using the StepMiner approach, all samples were sequentially ordered according to CRC progression (Normal → Adenoma → Cancer → Metastasis). Within this framework, each gene’s step index represented the precise point along the ordered progression. This approach integrates differential expression and progression patterns, enabling the identification of biologically meaningful progression.

**Table 2:** Top 10 Progression Genes.

| ProbeID | StepIndex | F_Stat | Direction | SYMBOL |
| --- | --- | --- | --- | --- |
| 211603_s_at | 74 | 234.6874359 | Up-Step | ETV4 |
| 203256_at | 73 | 218.9299516 | Up-Step | CDH3 |
| 204702_s_at | 74 | 208.8258437 | Up-Step | NFE2L3 |
| 218412_s_at | 74 | 202.096958 | Up-Step | GTF2IRD1 |
| 203908_at | 74 | 179.450834 | Down-Step | SLC4A4 |
| 205480_s_at | 74 | 163.1122177 | Down-Step | UGP2 |
| 202311_s_at | 117 | 157.6617675 | Up-Step | COL1A1 |
| 201479_at | 74 | 152.9993757 | Up-Step | DKC1 |
| 202242_at | 74 | 147.8722369 | Down-Step | TSPAN7 |
| 211494_s_at | 74 | 143.8467116 | Down-Step | SLC4A4 |

### 3.4 Biological Transition Mapping Across Colorectal Progression

Figure 3 demonstrates that the transition mapping adenoma-to-carcinoma phase has the highest number of major transcriptional shifts (StepIndex ≈74), suggesting this stage is a dominant molecular inflection point. Overall, 683 genes were involved in the normal-to-adenoma transition, 763 genes in the adenoma-to-primary cancer transition, and 608 genes in the cancer-to-metastatic transition. Both upregulated (Up-Step) and downregulated (Down-Step) genes were distributed across all biological transitions. These findings establish distinct stage-specific molecular checkpoints according to Table 3.

**Figure 3:**
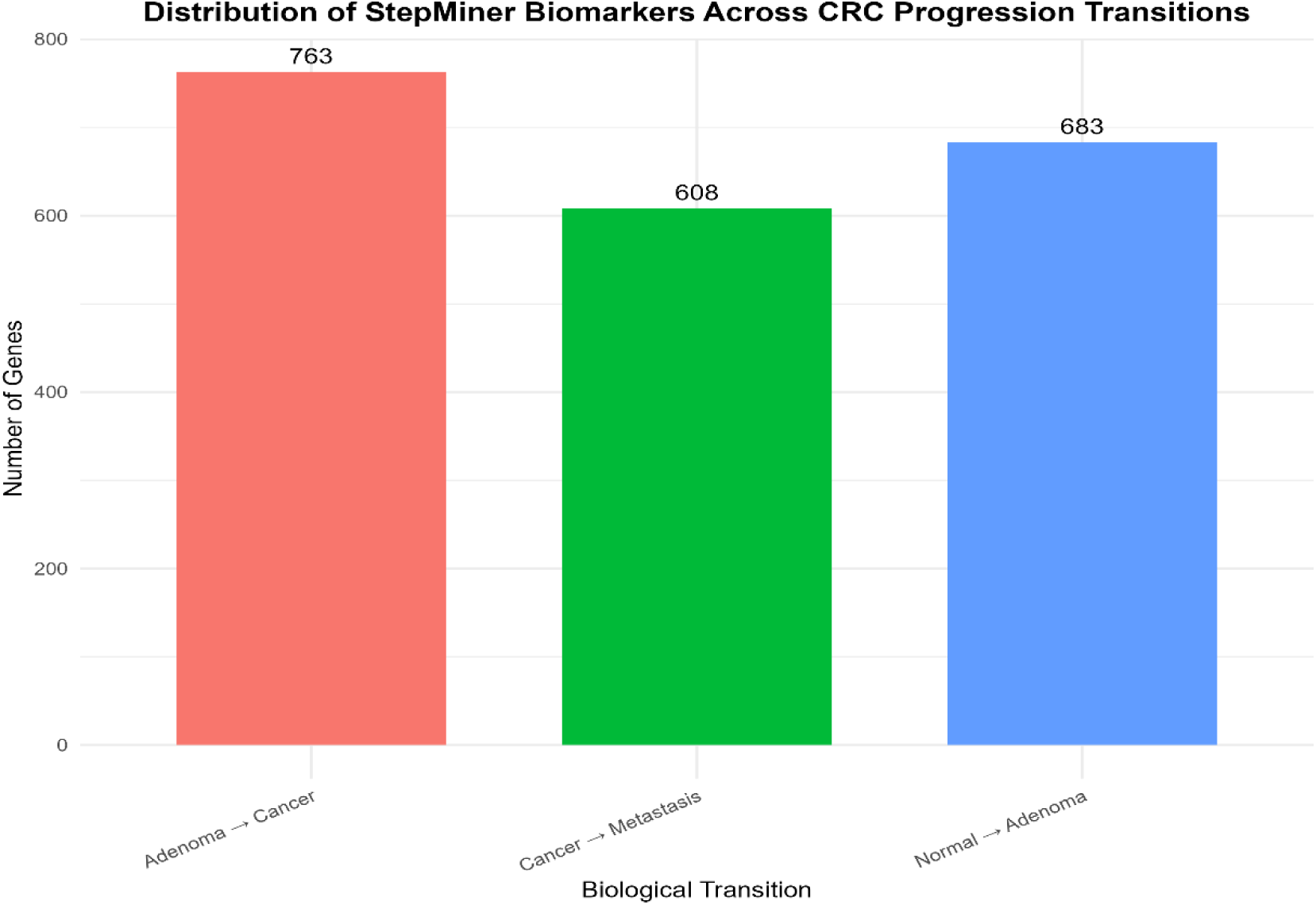
CRC Progression Transition.

**Table 3:** Progression Transition.

| Transition | Down-Step | Up-Step |
| --- | --- | --- |
| Adenoma → Colorectal Cancer | 355 | 408 |
| Normal Colon → Adenoma | 280 | 403 |
| Colorectal Cancer → Metastasis (Liver/Lung) | 272 | 336 |

### 3.5 State Classification Strategy

Balanced state-aware selection initially retained 25 top-ranked genes per transition. After removing overlapping genes across categories, a final set of 74 unique candidate genes was obtained. Selected 74 gene features. Using a 70/30 train-test split, the model achieved an OOB error rate of 15.97% and an overall accuracy of 88.5% in independent tests. Table 4 shows classification performance for normal colon (F1 = 0.980), adenoma (F1 = 0.875), and primary colorectal cancer (F1 = 0.898), while metastatic lesions showed lower sensitivity (0.571) but achieved high precision (0.923). Figure 4 displays the confusion matrix, which evaluates the model’s classification performance. The observed sensitivity likely reflects the documented clonality and molecular overlap between late-stage primary tumors and metastatic lesions [33]. That said, these metastatic cells still carry the transcriptional signature of the primary tumor. Identified COL1A1, VEGFA, NFE2L3, INHBA, and GTF2IRD1 in feature importance analysis, and those are major contributors to stage discrimination. Figure 5 illustrates the top 15 biomarkers in stage classification.

**Figure 4:**
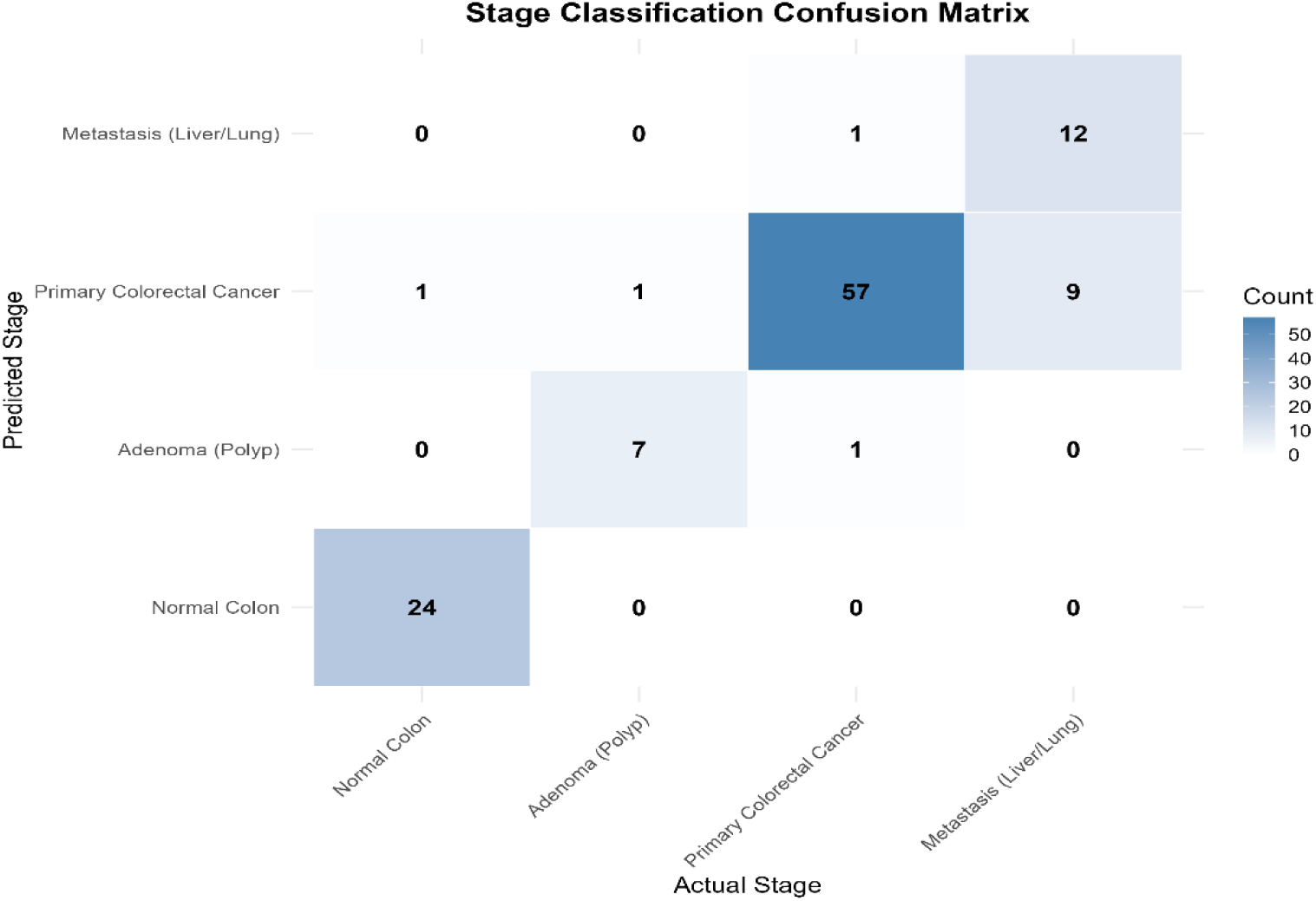
Stage Classification Confusion Matrix.

**Figure 5:**
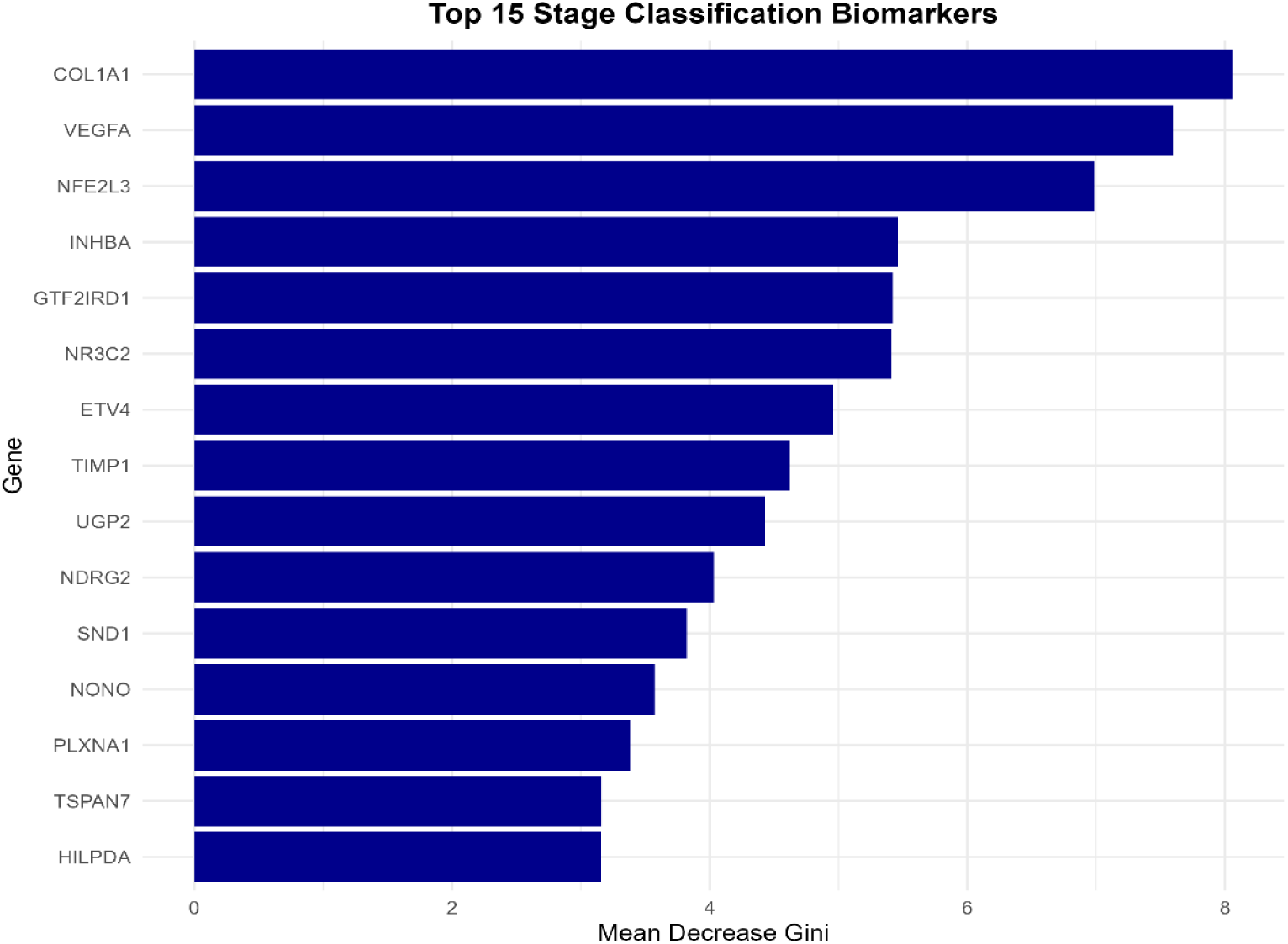
Top Stage Classification Biomarkers.

**Figure 6:**
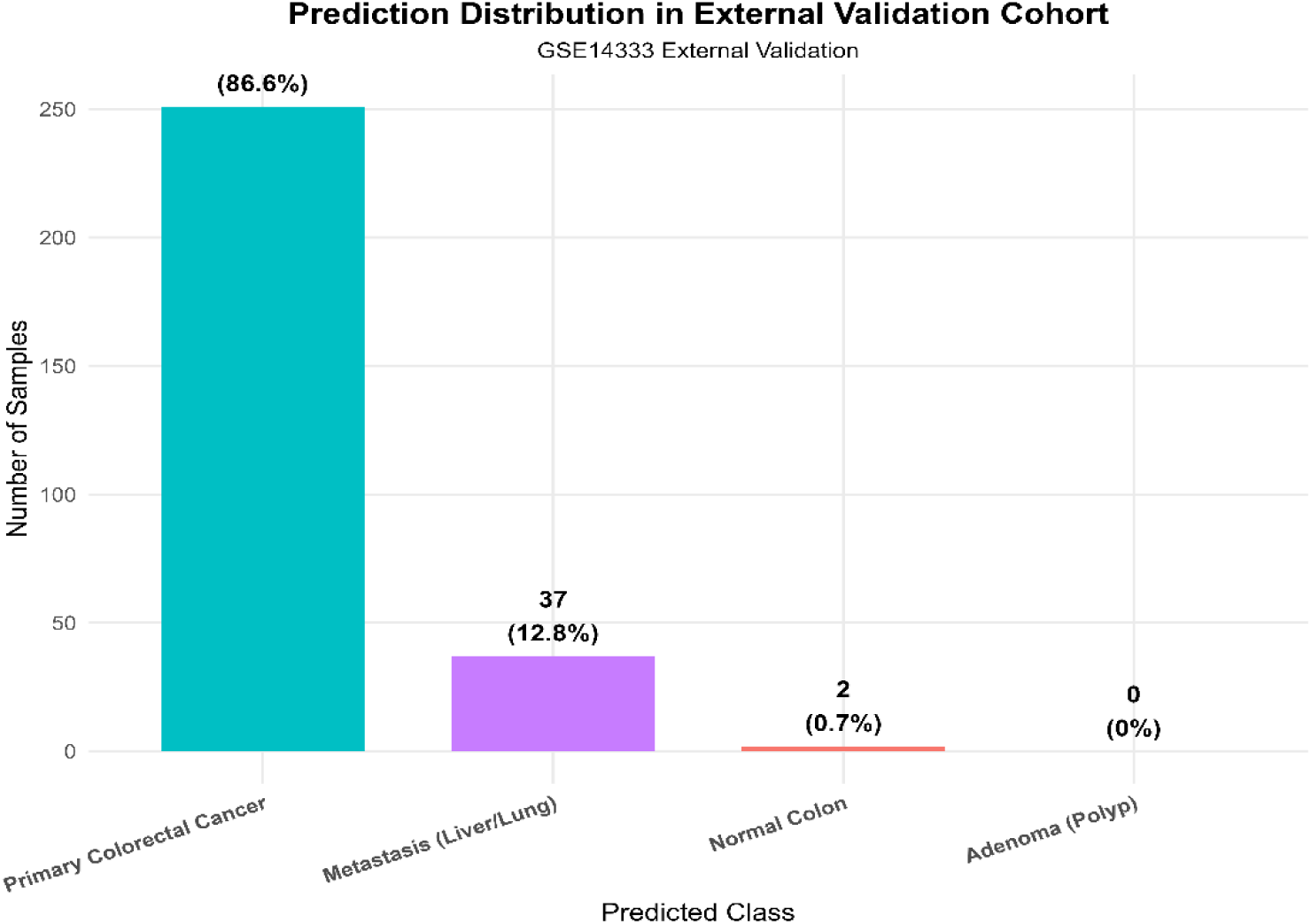
Prediction External Validation Cohort.

**Table 4:** Performance Per Class.

|  |  |  |  |
| --- | --- | --- | --- |
| Normal Colon | Sensitivity:<br>0.960 | Precision:<br>1.000 | F1:<br>0.980 |
| Adenoma (Polyp) | Sensitivity:<br>0.875 | Precision:<br>0.875 | F1:<br>0.875 |
| Colorectal Cancer | Sensitivity:<br>0.966 | Precision:<br>0.838 | F1:<br>0.898 |
| Metastasis (Liver/Lung) | Sensitivity:<br>0.571 | Precision:<br>0.923 | F1:<br>0.706 |

### 3.6 External Validation

To assess model generalizability, the trained model was applied to an independent GEO (GSE14333) consisting of primary colorectal cancer samples. After cross-platform probe harmonization and gene symbol mapping, Primary cancer prediction accuracy was 86.6%. A subset of samples was predicted as metastatic (n = 37), while only two samples were misclassified as normal tissue. These findings demonstrate the relevance of the progression-associated gene signature (Figure 2).

Also, we evaluated the trained model on an independent tissue-based cohort (GSE110223), achieving an AUC of 0.92. blood-based (GSE164191) and plasma-based (GSE142987) GEO datasets, performance dropped substantially (AUC ≈ 0.47), indicating random classification (Table 5). These findings indicate the biological differences between tissue and liquid-biopsy-based transcriptomic profiles, emphasizing the need for specific model optimization. Figure 7 shows a comparison curve between blood, tissue, and plasma.

**Figure 7:**
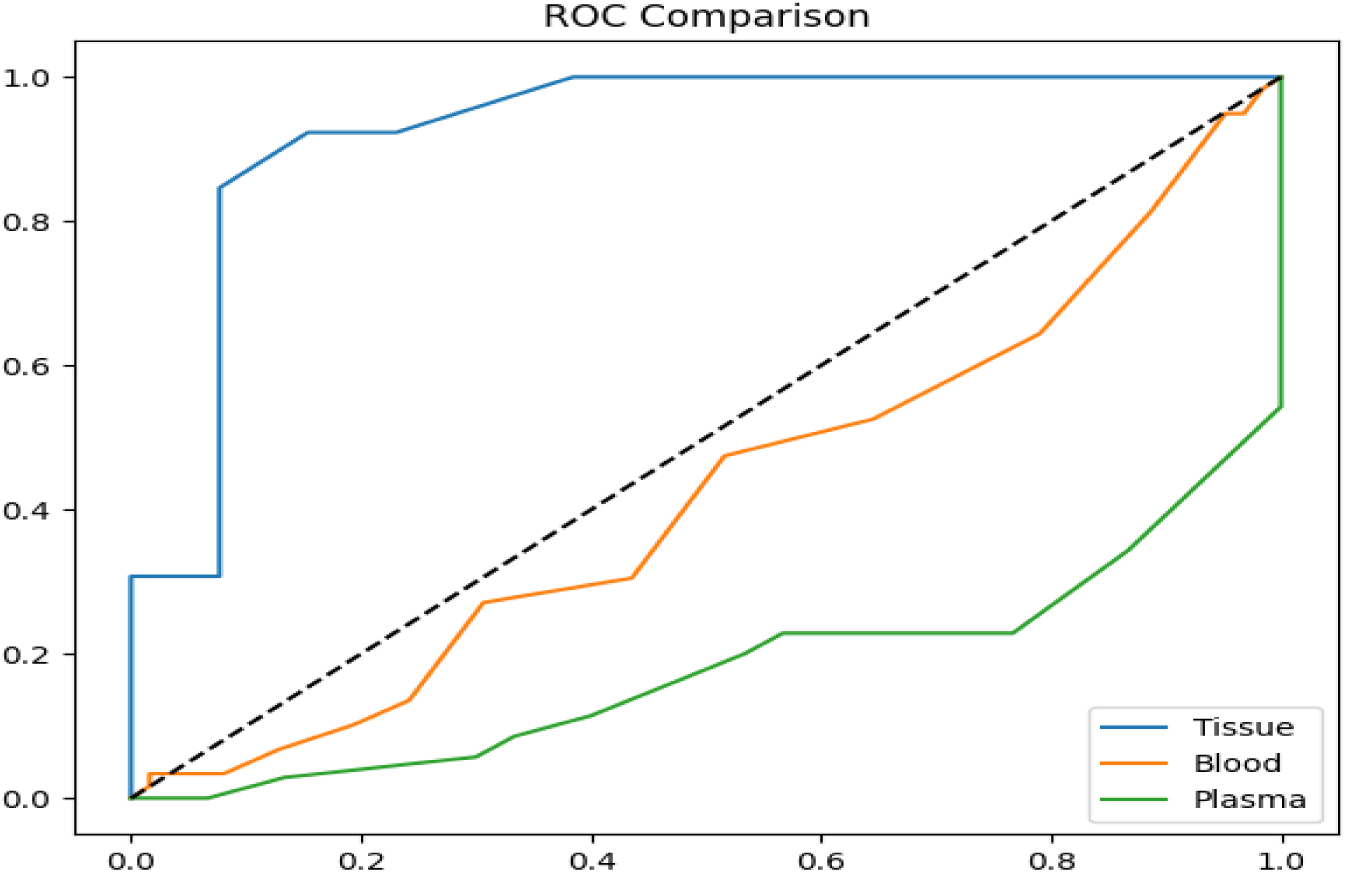
Validation in Tissue, Blood, and Plasma.

**Table 5:** External Validation Performance Table.

| GEO | AUC | Accuracy | Sensitivity | Specificity | F1-score |
| --- | --- | --- | --- | --- | --- |
| GSE110223 (Tissue) | 0.92 | 0.88 | 0.84 | 0.92 | 0.88 |
| GSE164191 (Blood) | 0.42 | 0.47 | 0.06 | 0.87 | 0.11 |
| GSE142987 (Plasma) | 0.20 | 0.46 | 0.0 | 1.0 | 0.0 |

### 3.7 Blood-Specific Model Refinement

To address poor blood performance, cross-filtering was performed to derive a refined gene set. Tissue-derived genes were filtered based on their relevance to blood expression (GSE164191). According to Figure 8, the refined model achieved a mean AUC of 0.868 ± 0.086, Accuracy: 0.818, Precision: 0.868, and F1-score: 0.803 on the GSE164191 blood dataset. This demonstrates that biologically informed gene filtering significantly improves model performance.

**Figure 8:**
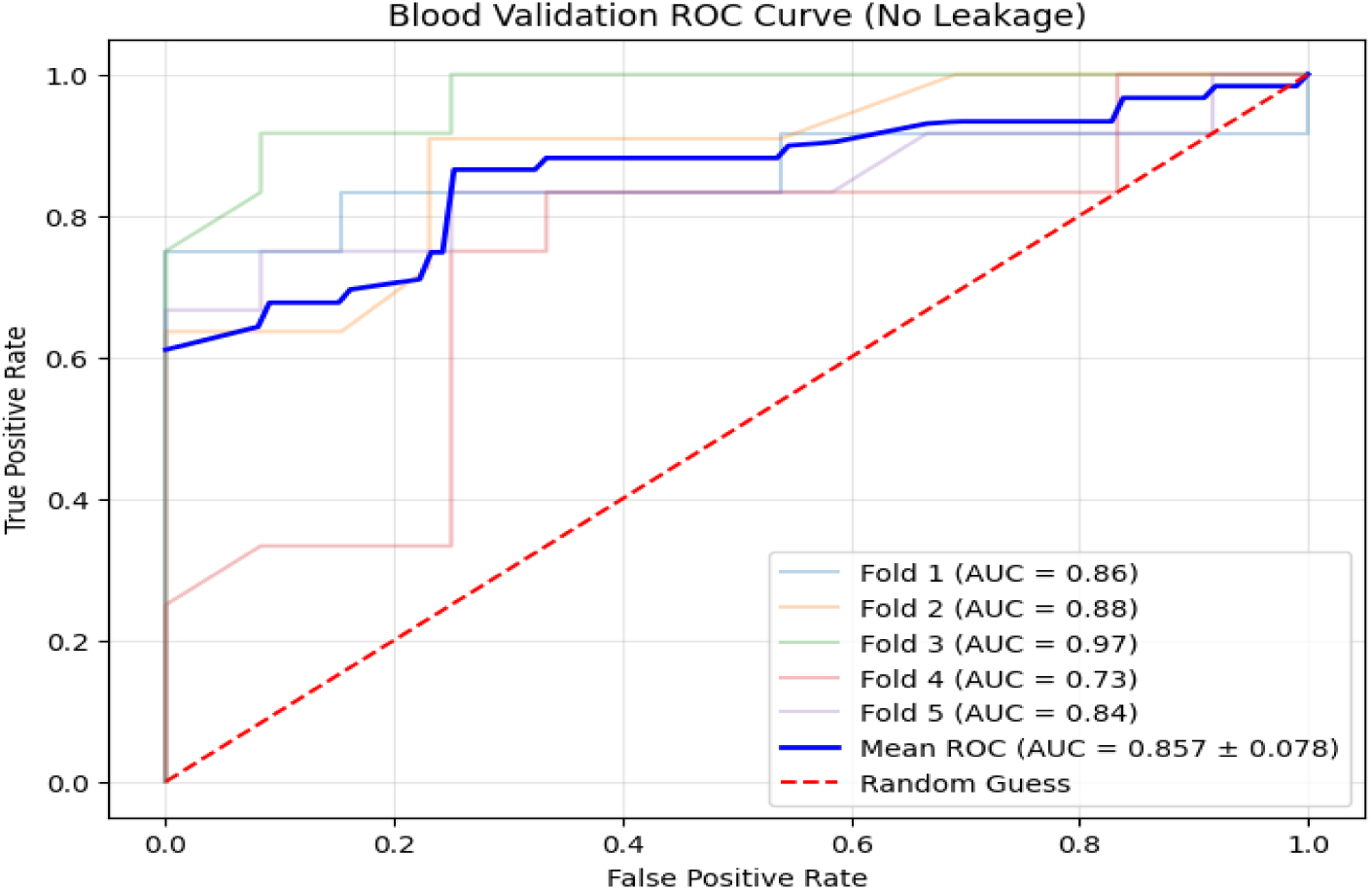
Gene Refinement Validation.

Although our initial signature contained 74 genes, our filtering identified 108 probes that remained significant (P < 0.05) in the blood. Multiple probes for genes such as CBX3, S100A11, and CCT6A showed consistent selection across all 5-fold cross-validation cycles (5/5 frequency), demonstrating their robustness as circulating biomarkers, as shown in Table 6.

**Table 6:** Gene After Filter In Blood.

| Probe_ID | Gene_Symbol | Selection_Frequency |
| --- | --- | --- |
| 1555920_at | CBX3 | 5 |
| 200660_at | S100A11 | 5 |
| 200037_s_at | CBX3 | 5 |
| 1562321_at | PDK4 | 5 |
| 200856_x_at | NCOR1 | 5 |
| 200816_s_at | PAFAH1B1 | 5 |
| 200815_s_at | PAFAH1B1 | 5 |
| 201416_at | SOX4 | 5 |
| 201327_s_at | CCT6A | 5 |

### 3.8 Functional Enhancement Analysis

Among the originally identified 74 tissue-derived colorectal cancer signature genes, 62 unique genes were successfully mapped. Functional enrichment analyses were performed using the blood-detectable genes. KEGG pathway enrichment analysis indicates involvement in cancer-associated pathways, nitrogen metabolism, AGE-RAGE signaling, pentose and glucuronate interconversion, starch and sucrose metabolism, and the PI3K-Akt signaling pathway (Figure 9).

**Figure 9:**
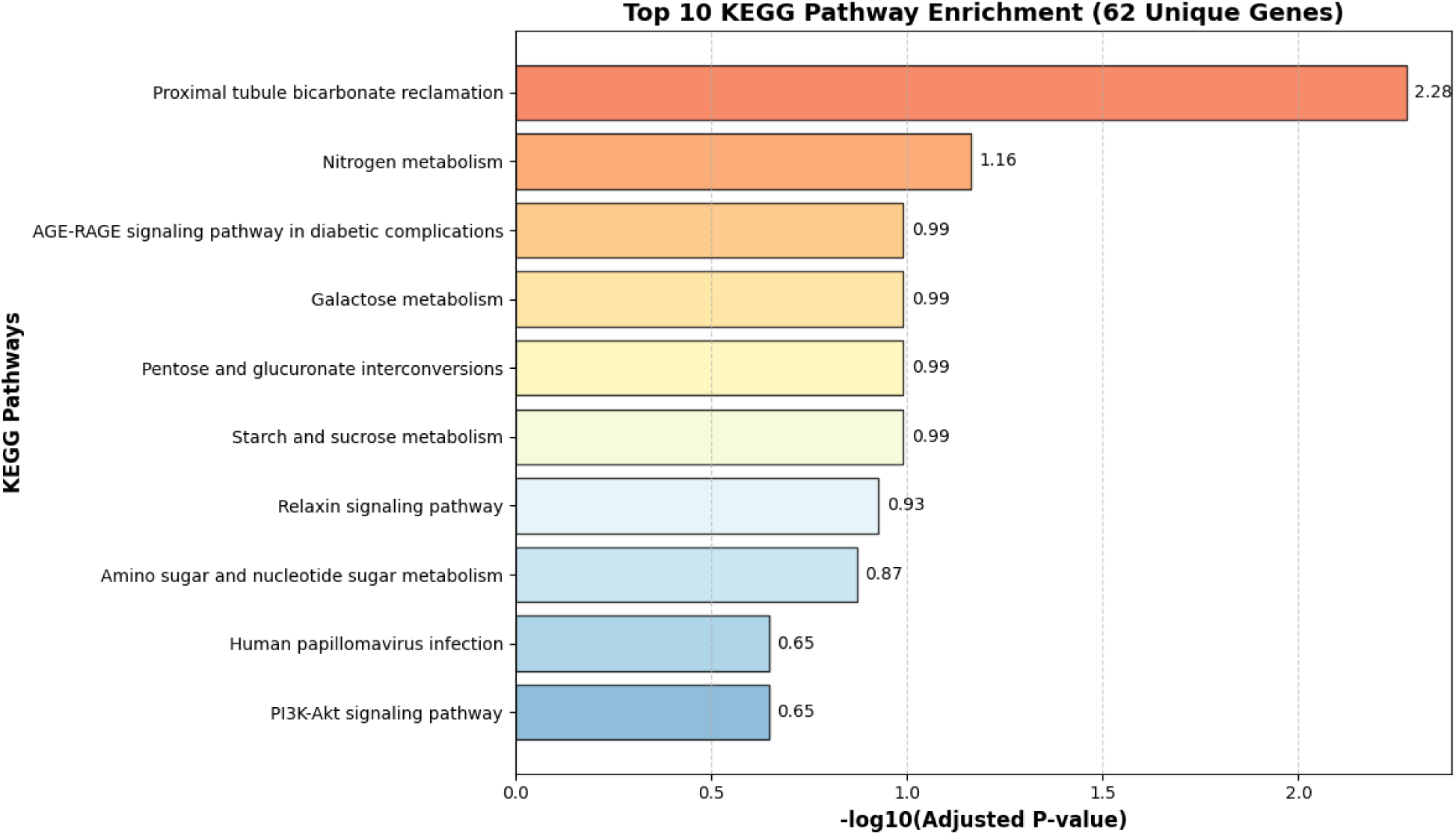
KEGG Pathway Analysis.

GO enrichment analysis revealed immune-related mechanisms, particularly neutrophil activation involved in immune response, neutrophil degranulation, bicarbonate transport, and protein localization-related mechanisms (Figure 10). These findings suggest that the identified genes may reflect both tumor-associated metabolic dysregulation and how the immune system reacts to the tumor in colorectal cancer.

**Figure 10:**
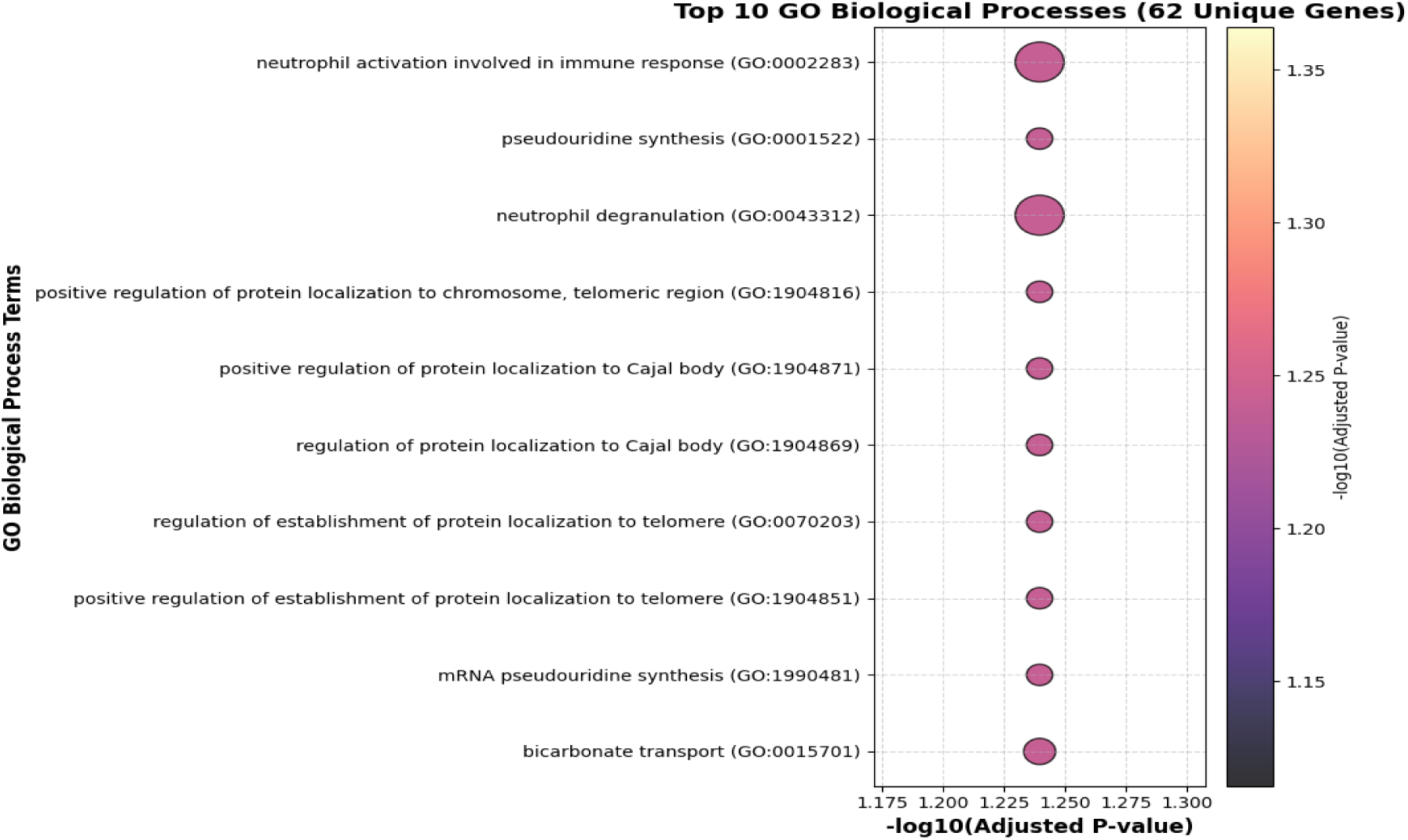
GO Pathway Analysis.

## 4 Discussion

We have developed a multi-stage (CRC) classification using transcriptomic signatures obtained from publicly available GEO datasets. This strategy was designed to show continuous changes in expression across the entire stages (Normal → Adenoma → CRC → Metastasis), rather than to analyze isolated stage comparisons. This focus on progression analysis allows us to identify the critical genetic drivers of colorectal cancer, rather than stage-specific biomarkers. By integrating analysis using StepMiner-inspired transition modeling and a monotonic approach, machine learning, and cross-platform external validation, we identified a set of meaningful, biologically important genes.

A major strength of this work lies in the procedure of selecting genes. Instead of relying solely on conventional differential expression analysis, we focused on genes showing a steady increase or decrease across disease stages (Normal → Adenoma → CRC → Metastasis), which helped us identify gene involvement in tumor progression. Then, a StepMiner-inspired transition-modeling approach was applied to identify genes that clearly switch between disease states. We used strict fold-wise feature selection to maintain rigor.

A Random Forest model achieved 86.9% accuracy on GSE14333 and an AUC of 0.9112 on GSE110223. Overall, our 74-gene signature performed better on external tissue datasets. However, performance dropped on the blood and plasma datasets (GSE164191, GSE142987), as expected, due to low tumor RNA levels and technical differences.

To address these differences, we refined our 74-gene signature to 62 unique genes by cross-filtering with blood data, and this new signature performed much better, achieving a mean AUC of 0.868. Including CBX3, S100A11, PDK4, NCOR1, PAFAH1B1, SOX4, and CCT6A. Many of these genes are known to be involved in colorectal tumorigenesis, metabolic adaptation, chromatin remodeling, and metastatic progression[34–37]. That connection validates that our proposal framework is biologically sound.

The functional enrichment results from our GO and KEGG analyses show that the 62 genes are associated with real biological processes in the body affected by colorectal cancer (CRC) in the blood.

Our GO analysis shows that many of these genes are involved in Neutrophil activation and degranulation. Neutrophils are the first immune cells to respond in the body when there is inflammation or cancer. Since we are using blood data, their activation and degranulation act like a "sensor" that detects the body’s primary response to a tumor[38,39].

KEGG analysis indicates metabolic pathways, such as nitrogen and Carbohydrate metabolism. We know how cancer cells change the body’s energy to help the tumor grow. Surprisingly, we also found pathways related to pH regulation (bicarbonate reclamation)[40,41]. This indicates that our gene signature can detect the body’s internal environment as cancer progresses. Lastly, the PI3K-Akt signaling pathway indicates that the gene signature is linked to major cancer drivers and acts as fuel for cell growth, survival, and anti-apoptotic signaling in colorectal cancer. [42]. In summary, these genes work together to produce a multidimensional biomarker that captures how the body responds to colorectal cancer, through both the immune system and metabolism.

However, we acknowledge several limitations. First, Internal validation was performed using the GSE41258 dataset, which includes normal, adenoma, cancer, and metastasis samples. External and cross-validation datasets contained only colorectal cancer and normal samples. Second, we are not evaluating multi-omics data such as genomics, proteomics, or epigenomics, which may provide more biological insights.

Also, we did not perform experimental validation at the clinical level, and multi-omics functional studies are warranted before clinical use.

## 5 Conclusion and Future Work

Overall, this study highlights that our method of combining progression analysis and a transition approach works well for identifying meaningful CRC biomarkers. Also clearly reveals the real challenge of using tissue biomarkers in blood or plasma samples. Although the tissue classifier showed strong external reproducibility.

This 62-gene signature offers a robust approach for detecting colorectal cancer from blood samples. Our functional analysis using GO and KEGG confirms that these genes are associated with the biological reality of CRC, specifically highlighting the roles of immune activation and metabolic adaptation [37,39,42]. Our aim is to validate these findings in larger clinical cohorts and explore the development of a simplified PCR-based diagnostic kit. By bridging the gap between computational feature selection and biological mechanisms, this work provides a good foundation for CRC detection and improved patient outcomes in clinical practice.

## 6 Data and Code Availability

We obtained all gene expression data from the NCBI Gene Expression Omnibus (GEO) database. Our application’s source code (Angular, ASP.NET Core, and R Plumber) is publicly available on GitHub at the [link](https://github.com/Pritom300/CRC-APP). The reproducible code behind the workflow is available from the corresponding author upon request.

## Reference

1. Keum N, Giovannucci E. Global burden of colorectal cancer: emerging trends, risk factors and prevention strategies. Nat Rev Gastroenterol Hepatol. 2019;16: 713–732. doi:10.1038/s41575-019-0189-8

2. Xi Y, Xu P. Global colorectal cancer burden in 2020 and projections to 2040. Transl Oncol. 2021;14: 101174. doi:10.1016/j.tranon.2021.101174

3. Sung H, Ferlay J, Siegel RL, Laversanne M, Soerjomataram I, Jemal A, et al. Global Cancer Statistics 2020: GLOBOCAN Estimates of Incidence and Mortality Worldwide for 36 Cancers in 185 Countries. CA Cancer J Clin. 2021;71: 209–249. doi:10.3322/caac.21660

4. Arnold M, Sierra MS, Laversanne M, Soerjomataram I, Jemal A, Bray F. Global patterns and trends in colorectal cancer incidence and mortality. Gut. 2017;66: 683–691. doi:10.1136/gutjnl-2015-310912

5. Zhou L, Yu L, Liao M, Peng T, Zhang L, Han C, et al. Integrating machine learning and genetic evidence to uncover novel gene biomarkers for colorectal cancer diagnosis. Discov Oncol. 2025;16: 675. doi:10.1007/s12672-025-02435-0

6. Yin Y, Yang Z, Li X, Gong S, Xu C. Gene Expression-Based Colorectal Cancer Prediction Using Machine Learning and SHAP Analysis. Genes. 2026;17: 114. doi:10.3390/genes17010114

7. Vaziri-Moghadam A, Foroughmand-Araabi M-H. Integrating machine learning and bioinformatics approaches for identifying novel diagnostic gene biomarkers in colorectal cancer. Sci Rep. 2024;14: 24786. doi:10.1038/s41598-024-75438-6

8. Wei W, Li Y, Huang T. Using Machine Learning Methods to Study Colorectal Cancer Tumor Micro-Environment and Its Biomarkers. Int J Mol Sci. 2023;24: 11133. doi:10.3390/ijms241311133

9. Tsang HF, Pei XM, Wong YKE, Wong SCC. Plasma Circulating mRNA Profile for the Non-Invasive Diagnosis of Colorectal Cancer Using NanoString Technologies. Int J Mol Sci. 2024;25: 3012. doi:10.3390/ijms25053012

10. Kaya IH, Al-Harazi O, Colak D. Transcriptomic data analysis coupled with copy number aberrations reveals a blood-based 17-gene signature for diagnosis and prognosis of patients with colorectal cancer. Front Genet. 2023;13. doi:10.3389/fgene.2022.1031086

11. Alawad DM, Fertel M, Hicks C. Leveraging Multi-Model Machine Learning Algorithms for Tumor–Normal Classification and Discovery of Biomarkers in Colorectal Cancer Using Multi-Omics Data. Cancers. 2026;18: 1503. doi:10.3390/cancers18101503

12. Yin Y, Yang Z, Li X, Gong S, Xu C. Gene Expression-Based Colorectal Cancer Prediction Using Machine Learning and SHAP Analysis. Genes. 2026;17: 114. doi:10.3390/genes17010114

13. Alawad DM, Fertel M, Hicks C. Leveraging Multi-Model Machine Learning Algorithms for Tumor–Normal Classification and Discovery of Biomarkers in Colorectal Cancer Using Multi-Omics Data. Cancers. 2026;18: 1503. doi:10.3390/cancers18101503

14. Gallitto G, Englert R, Kincses B, Kotikalapudi R, Li J, Hoffschlag K, et al. External validation of machine learning models-registered models and adaptive sample splitting. GigaScience. 2025;14: giaf036. doi:10.1093/gigascience/giaf036

15. Li H, Qin J, Li Z, Ouyang R, Chen Z, Huang S, et al. Systematic review and meta-analysis of deep learning for MSI-H in colorectal cancer whole slide images. Npj Digit Med. 2025;8: 456. doi:10.1038/s41746-025-01848-z

16. Yuan T, Edelmann D, Kather JN, Fan Z, Tagscherer KE, Roth W, et al. DNA methylation-based biomarkers and prediction models for the survival of patients with colorectal cancer: systematic review and external validation study. medRxiv; 2022. p. 2022.11.03.22281595. doi:10.1101/2022.11.03.22281595

17. Sahoo D, Dill D, Tibshirani R, Plevritis S. Extracting binary signals from microarray time-course data. Nucleic Acids Res. 2007;35: 3705–12. doi:10.1093/nar/gkm284

18. Identification of diagnostic biomarkers via weighted correlation network analysis in colorectal cancer using a system biology approach | Scientific Reports. [cited 31 May 2026]. Available: https://www.nature.com/articles/s41598-023-40953-5

19. Ashburner M, Ball CA, Blake JA, Botstein D, Butler H, Cherry JM, et al. Gene Ontology: tool for the unification of biology. Nat Genet. 2000;25: 25–29. doi:10.1038/75556

20. Kanehisa M, Goto S. KEGG: Kyoto Encyclopedia of Genes and Genomes. Nucleic Acids Res. 2000;28: 27–30. doi:10.1093/nar/28.1.27

21. GEO Accession viewer. [cited 13 May 2026]. Available: https://www.ncbi.nlm.nih.gov/geo/query/acc.cgi?acc=GSE41258

22. Qiu C, Zhang Y, Chen L. Impaired Metabolic Pathways Related to Colorectal Cancer Progression and Therapeutic Implications. Iran J Public Health. 2020;49: 56–67.

23. Ke T-W, Chang S-C, Yeh C-M, Lin S-H, Yeh K-T. Comprehensive bioinformatic analysis reveals prognostic significance and functional insights of candidate gene expression in colorectal cancer. Sci Rep. 2025;15: 5659. doi:10.1038/s41598-025-90025-z

24. GEO Accession viewer. [cited 13 May 2026]. Available: https://www.ncbi.nlm.nih.gov/geo/query/acc.cgi?acc=GPL570

25. Breiman L. Random Forests. Mach Learn. 2001;45: 5–32. doi:10.1023/A:1010933404324

26. (PDF) Scikit-learn: Machine Learning in Python. [cited 13 May 2026]. Available: https://www.researchgate.net/publication/51969319_Scikit-learn_Machine_Learning_in_Python

27. GEO Accession viewer. [cited 13 May 2026]. Available: https://www.ncbi.nlm.nih.gov/geo/query/acc.cgi?acc=GSE164191

28. GEO Accession viewer. [cited 13 May 2026]. Available: https://www.ncbi.nlm.nih.gov/geo/query/acc.cgi?acc=GSE110223

29. GEO Accession viewer. [cited 13 May 2026]. Available: https://www.ncbi.nlm.nih.gov/geo/query/acc.cgi?acc=GSE142987

30. GEO Accession viewer. [cited 13 May 2026]. Available: https://www.ncbi.nlm.nih.gov/geo/query/acc.cgi?acc=GSE14333

31. (PDF) Stratified K-fold cross validation optimization on machine learning for prediction. ResearchGate. [cited 13 May 2026]. doi:10.33395/sinkron.v7i4.11792

32. Kaliappan J, Bagepalli AR, Almal S, Mishra R, Hu Y-C, Srinivasan K. Impact of Cross-Validation on Machine Learning Models for Early Detection of Intrauterine Fetal Demise. Diagnostics. 2023;13: 1692. doi:10.3390/diagnostics13101692

33. Dang HX, Krasnick BA, White BS, Grossman JG, Strand MS, Zhang J, et al. The clonal evolution of metastatic colorectal cancer. Sci Adv. 2020;6: eaay9691. doi:10.1126/sciadv.aay9691

34. Fan Y, Li H, Liang X, Xiang Z. CBX3 promotes colon cancer cell proliferation by CDK6 kinase-independent function during cell cycle. Oncotarget. 2017;8: 19934–19946. doi:10.18632/oncotarget.15253

35. Leclerc D, Pham DNT, Lévesque N, Truongcao M, Foulkes WD, Sapienza C, et al. Oncogenic role of PDK4 in human colon cancer cells. Br J Cancer. 2017;116: 930–936. doi:10.1038/bjc.2017.38

36. Huang Y, Tang X, Xie H, Wu Z, Jin L, Zhang L, et al. USP14/S100A11 axis promote colorectal cancer progression by inhibiting cell senescence. Cell Death Dis. 2025;16: 384. doi:10.1038/s41419-025-07724-8

37. Wang B, Li Y, Tan F, Xiao Z. Increased expression of SOX4 is associated with colorectal cancer progression. Tumour Biol J Int Soc Oncodevelopmental Biol Med. 2016;37: 9131–9137. doi:10.1007/s13277-015-4756-5

38. Bai M, Jin Y, Jin Z, Xie Y, Chen J, Zhong Q, et al. Distinct immunophenotypic profiles and neutrophil heterogeneity in colorectal cancer. Cancer Lett. 2025;616: 217570. doi:10.1016/j.canlet.2025.217570

39. Yang Y, Lu C, Li L, Zheng C, Wang Y, Chen J, et al. Construction and multicohort validation of a colon cancer prognostic risk score system based on big data of neutrophil-associated differentially expressed genes. J Cancer. 2024;15: 2866–2879. doi:10.7150/jca.94560

40. Pang B, Wu H. Metabolic reprogramming in colorectal cancer: a review of aerobic glycolysis and its therapeutic implications for targeted treatment strategies. Cell Death Discov. 2025;11: 321. doi:10.1038/s41420-025-02623-5

41. Hozhabri H, Lashkari A, Razavi S-M, Mohammadian A. Integration of gene expression data identifies key genes and pathways in colorectal cancer. Med Oncol. 2021;38: 7. doi:10.1007/s12032-020-01448-9

42. Maharati A, Moghbeli M. PI3K/AKT signaling pathway as a critical regulator of epithelial-mesenchymal transition in colorectal tumor cells. Cell Commun Signal CCS. 2023;21: 201. doi:10.1186/s12964-023-01225-x

